# Oxygen production as an electron overflow pathway in ammonia-oxidizing archaea

**DOI:** 10.64898/2025.12.03.692013

**Authors:** Thomas Pribasnig, Christa Schleper, Logan H. Hodgskiss

## Abstract

Ammonia-oxidizing archaea (AOA) are widespread in nature and contribute to global carbon and nitrogen cycling. Although their energy production is an aerobic process, AOA occur in environments of low oxygen and in fully anoxic zones. Recently, oxygen production under anoxic conditions was demonstrated, raising the hypothesis that this activity can sustain oxygen dependent ammonia oxidation. In the work presented here, *Nitrososphaera viennensis* produces high amounts of oxygen in anoxia in dependence of nitrite but not ammonia. Rather than conferring a physiological benefit, oxygen production impaired recovery from anoxia. Produced oxygen is mostly not consumed and therefore cannot sustain ammonia oxidation in the absence of external oxygen. Instead, it is tightly linked to the amount of available redox equivalents. These observations indicate that oxygen production represents a general physiological strategy for alleviating the potential accumulation of reactive intermediates in the ammonia oxidation pathway, analogous to nitrifier denitrification in ammonia-oxidizing bacteria.

**Teaser:** Oxygen production in ammonia-oxidizing archaea is not an adaptation to anoxia, but rather an electron overflow pathway that is detectable in anoxic conditions.

## Intro

Ammonia-oxidizing archaea (AOA) gain energy by oxidizing ammonia to nitrite^1,2^, through which they contribute to the global nitrogen cycle^3,4^. AOA are remarkably efficient in their central metabolism. They employ the most energy efficient aerobic carbon fixation cycle^5^, and a type *aa3* terminal oxidase as Complex IV, the most efficient terminal oxidase in terms of protons pumped per electron^6,7^. The ammonia monooxygenase (AMO) requires input of 2 electrons to carry out the oxidation of NH_3_ to NH_2_OH^8–10^, which is further oxidized to NO_2_^−^ by one or more unknown enzymes for energy conservation^11–14^. AOA do not only have a remarkably high affinity for NH_3_^15–17^,but also have been shown to have a high affinity for oxygen, allowing activity under low oxygen concentrations^18^. It may be their extremely efficient metabolism that enables their ubiquitous distribution in almost every aerobic environment^19–21^. Unexpectedly, they have also been found to be highly abundant in environments with low or no oxygen^22–26^. Their occurrence in those environments is surprising, since AMO and the terminal oxidase both rely on oxygen for activity, making them dependent on oxygen for their metabolism.

The capability of *Nitrosopumilus maritimus*, and other AOA, to produce oxygen under anoxic conditions drastically changed that view^27,28^. In this newly proposed pathway, NO_2_^−^ is first reduced to NO, presumably by the nitrite reductase NirK, and subsequently dismutated to N_2_O and O_2_. Independent from this novel pathway, N_2_O production by AOA has been demonstrated in several studies, either as a byproduct of hydroxylamine oxidation or as a result of NO_2_^−^ reduction^18,29–32^. Under anoxic conditions, the reduction of N_2_O to N_2_ is species-specific^28,33^. While some strains, such as *N. viennensis,* stop at N_2_O, others, like *N. maritimus,* further reduce it to N_2_. Even the reduction of exogenous N_2_O has been described, making specific strains of AOA potential sinks for N_2_O under anoxic conditions^33^. Oxygen production could expand the range of AOA activity into anoxic environments. However, oxygen production requires more electrons than can be obtained from ammonia oxidation alone^27^. So far, all isolated AOA that have been shown to produce oxygen are strict autotrophs, and have not been shown to gain electrons from any other source than ammonia oxidation^1,30,34,35^. Why N_2_O would further be reduced to N_2_ is not compatible with the current interpretation of oxygen production to sustain activity under anoxic conditions. A wasteful pathway like this is in stark contrast to the otherwise extremely efficient metabolism employed by AOA and the need to streamline expenses even further if activity should be sustained under anoxic conditions.

In this study a range of physiological experiments was used to investigate the dependence of oxygen production in the soil ammonia oxidizer, *Nitrososphaera viennensis*. Oxygen production is proposed as a NirK dependent electron overflow pathway rather than an adaptation to anoxic conditions. It is shown that produced oxygen cannot sustain ammonia oxidation in the absence of external oxygen and is instead tightly linked to the amount of available redox equivalents. Taken together, these findings reshape the view of oxygen production in AOA, positioning it as a general physiological strategy for addressing the potential buildup of reactive intermediates in the ammonia oxidation pathway rather than an expansion of their ecological niche.

## Material and methods

### General cultivation and preparation of anoxic medium

#### Cultivation

*N. viennensis* was cultivated as previously described^30^. Basic freshwater medium (FWM) was supplemented with 2 mM NH_4_Cl, 2 mM NaHCO_3_, 7.5 µM FeNaEDTA, 1mM pyruvate, 50 µg/ml kanamycin, non-chelated trace elements (0.1% volume/volume (v/v)), and buffered with 10 mM HEPES to pH 7.5. All solutions were either sterilized by filtration (0.2 µm mixed cellulose ester (MCE) membrane filters), or autoclaving (FWM). Nitrite production was followed by colorimetric measurements via the Griess reaction as described previously. Cultures were incubated at 42°C, shaking at 80 rpm, in the dark. Cell numbers were determined as previously described, by linking NO_2_^−^ values to cell counts (Pribasnig, Dreer et al. in preparation). Scripts are available under: https://github.com/pribasnig/oxygen_production.

For experiments, cells were concentrated by centrifugation in a Sorvall Lynx 400 centrifuge using a Thermo Scientific Fiberlite F12-6x500 LEX rotor and 400 ml centrifugation vessels (21,000 x g, 30 min, 40°C), washed in NH_4_Cl-free growth medium to remove residual NH_4_Cl or NO_2_^−^ and centrifuged again. Cell pellets were resuspended in a known volume of NH_4_Cl-free growth medium, pooled and distributed into the respective experimental conditions. The amounts of cells used were similar over all experiments (7.0*10^9^±6.8*10^8^ cells, DatasetS1). Cell specific oxygen production rates (amol*cell^-1^) were calculated by dividing net production (µmol) by cell numbers (DatasetS1).

#### Anaerobic preparations

120 ml Serum bottles were equipped with Presense PSt6-YAU-D5-YOP oxygen sensor spots, following the manufacturer’s instructions, autoclaved twice, and filled with 80 ml of respective medium under sterile conditions. Bottles were sealed airtight using 20 mm blue butyl rubber stoppers and 20 mm aluminium crimp caps to seal the bottles, then sparged with N_2_ for 30 min, incubated overnight at 42°C (80 rpm), and sparged again for 30 min. Oxygen concentrations were measured with a Fibox4 Optode. If concentrations were > 0.5 µM, flasks were sparged further until concentrations ≤ 0.5 m µM were reached. Near-anoxic flasks were transferred to an anoxic chamber, where they were completely filled with-near anoxic medium prepared the same way, using sterile syringes and needles, leaving only a small N2 bubble for incubations later. Bottles were transferred to 42°C for at least 24 h before experiments were started and O_2_ concentrations measured again. All conditions contain 2 mM NH_4_^+^ and 1 mM NO_2_^−^ unless otherwise specified.

#### Incubations with different amounts of NO_2_^−^ and NH_4_^+^

To test the dependence on NO_2_^−^, near-anoxic medium was prepared with either 1 mM, 0.05 mM, 10 mM, or 0 mM NO_2_^−^. To assess the influence of NH_4_^+^, near-anoxic medium was prepared with either 2 mM or 0 mM NH_4_Cl. In each treatment 0.5 ml of concentrated cells were inoculated into the medium and maintained in a waterbath at 42°C, and oxygen consumption and production monitored over 24 h. Cell numbers for each experiment can be found in DatasetS1.

#### Recovery after anoxic conditions

After 24 h of anoxic incubation, cultures either oxygen producing (with NO _2_^−^) or not (without NO_2_^−^), were returned to optimal oxic growth conditions. For this, bottles were opened sterilely and 1 ml (6.12*10^7^ cells) was inoculated immediately into 20 ml of 42°C prewarmed, optimal growth medium in 30 ml Greiner tubes. Growth was followed over time via nitrite measurements as described above.

#### Starvation experiments

Cells were resuspended and pooled as described above, then divided into three groups. The first group was immediately inoculated into near-anoxic medium containing 1 mM NO_2_^−^ and 2 mM NH_4_Cl. The second and third groups were incubated in 20 mL of NH_4_Cl-free medium at 42 °C for 24 h or 72 h, respectively, to starve cells. After those incubations, starved cells were centrifuged, resuspended, and inoculated into near-anoxic medium containing 1 mM NO_2_^−^. and 2 mM NH_4_Cl. Oxygen production was monitored over 24 hours in all treatments.

#### Oxygen additions

To test the effect of oxygen additions, cells were inoculated into near-anoxic medium containing 1 mM NO_2_^−^ under three conditions: Condition a) with 2 mM of NH_4_Cl, Condition b) with 0 mM NH_4_Cl and Condition c) with 2 mM of NH_4_Cl and 3 mM of allylthiourea (ATU). In Condition a) replicates were split into 2 groups. When oxygen was produced to a concentration of ∼1 µM, the first group received an addition of 0.5 ml 42°C oxic growth medium (containing 98.5 µmol oxygen), leading to an O_2_ increase of 0.86 µM, and the second group received 1 ml (containing 197µmol oxygen), leading to an O_2_ increase of 1.73 µM. For the second group, the same additions (containing 197 µmol oxygen), leading to an O_2_ increase of 1.73 µM each time, were repeated twice more, at concentrations of 1 µM and 1.4 µM.

Condition b) replicates were split in 2 groups. No additions were made to the first group. The second group received 2 additions of 1 ml 42°C oxic growth medium (containing 197 µmol oxygen) leading to an increase of 1.73 µM for each addition. Additions were made when oxygen was produced to concentrations of ∼1 µM.

Condition c) replicates were split into 2 groups. After oxygen was produced to a concentration of ∼0.8 µM, one group received an addition of 1 ml 42°C anoxic medium, the other group 1 ml 42°C anoxic medium containing 345 mM ATU, resulting in a final concentration of 3 mM ATU.

Additions of anoxic medium alone did not influence oxygen production. When oxygen was produced to a concentration of ∼1.5 µM, both groups received additions of 1 ml 42°C oxic growth medium (containing 197 µmol oxygen), increasing concentrations by 1.73 µM. In all experiments oxygen consumption and production was measured over 24 hours.

### Transcriptome analysis

To conduct transcriptomic studies, *N. viennensis* was grown in 500 mL optimal oxic medium. Upon reaching mid-exponential phase, cultures were either directly filtered to use as a control, or centrifuged and inoculated in near-anoxic medium, where oxygen production was monitored. To harvest cultures after 2 h or 24 h under oxygen producing conditions, cultures were opened and filtered in an anoxic tent, filters stored in Falcon tubes, flash frozen and incubated at −70°C. RNA was extracted using the mirVana miRNA Isolation Kit (Thermo Fisher) according to the manufacturer’s instructions and as described before^36^. RNA was submitted to the Vienna Biocenter Facility (VBC) for sequencing with a species specific rRNA depletion step. Samples from the 24 h experiment were sequenced as 100 base pair singleend reads and samples from the 2 h experiment were sequenced as 150 base pair paired-end reads. Raw reads were downloaded from VBC, checked with md5sum and are available at the NCBI repository under BioProject ID: PRJNA1358245. FastQC v.0.12.1 and multiQC v.1.25 were used to analyze the reads. Reads were trimmed using fastp v.0.23.4 with specific settings: rim_front1 12 -- trim_front2 12 --trim_poly_g --detect_adapter_for_pe --average_qual 30 --length_required 30. SortmeRNA v.4.3.4 was used to sort out rRNA reads, and identified mRNA reads were mapped to the genome using HISAT2 v.2.2.1 and counted using featureCounts v2.1.1. To calculate TPM values, the read counts were divided by the length of each gene in kilobases, which yielded reads per kilobase (RPK). The sum of all RPK values was divided by 1000000 giving a per-million scaling factor, which was used to divide individual RPK values to yield transcripts per million (TPM). Since the anoxic transcriptomes were degraded, differential expression analysis was not carried out.

### Statistical methods

Total produced oxygen data was plotted for each condition after 24 hours (Dataset S1). Data was tested in R version 4.5.1^37^ using Rstudio 2025.09.1.401^38^ for normality and homogeneity of variance using the Shapiro test (shapiro_test, car package v3.1-3^39^) and Levene test (leveneTest, rstatix v0.7.3 package^40^) respectively. While all conditions passed the normality check, the assumptions for homogeneity of variance were not met. Therefore, conditions were evaluated for differences in means using the Games-Howell test (GamesHowellTest, PMCMRplus v1.9.12 package^41^) which is suitable for data with unequal variances. Post-hoc analysis (posthocTGH, rosetta v0.3.12 package^42^) was used to obtain adjusted p-values using the Benjamini-Hochberg method and conditions were placed into statistically similar groups using the multcompLetters() function of the multcompView v0.1-10 package^43^. The R package tidyverse v2.0.08 was used for helpful processing of data^44^.

## Results

### Oxygen production is dependent on NO_2_^−^ and independent from NH_4_^+^

*N. viennensis* cells were washed to remove residual NO_2_^−^ and NH_4_^+^ and inoculated into a near-anoxic medium with or without NO_2_^−^ (Fig. 1a) or NH_4_^+^ (Fig. 1c) to initial concentrations of 6.12*107 cells/ml. In both conditions, the introduced oxygen was rapidly consumed to ∼0.2- 0.3 µM. In the presence of NO_2_^−^, oxygen production started immediately after and continued for 24 h, with the highest rates in the first hours and a gradual slowdown over time. The total oxygen concentration measured (3.5 µM) was an order of magnitude higher than reported in previous studies, due to the use of concentrated cells (Fig. 1a). In the absence of NO_2_^−^, no increase in oxygen levels was observed after the initial depletion and oxygen levels stayed constant at ∼0.3 µM over 24 h (Fig. 1a). Experiments with higher nitrite resulted in larger concentrations of oxygen, reinforcing the dependence on nitrite (Fig. 1b).

**Figure 1:**
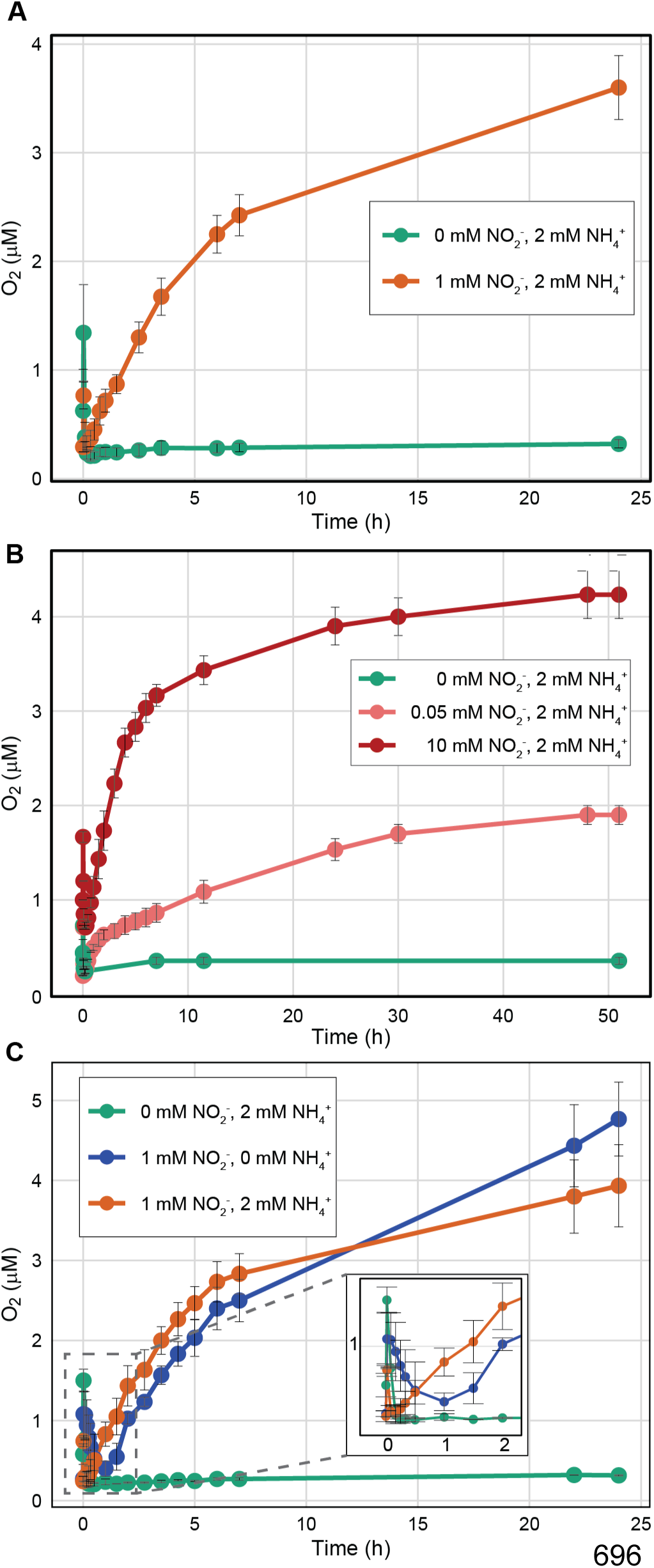
Oxygen production is dependent on NO2^−^ and independent from NH4^+^ : Oxygen production (in µM) of *N. viennensis* under anoxic conditions over time (in h). Conditions contain 2 mM NH_4_ ^+^ and 1 mM NO_2_^−^ unless otherwise specified. **(A)** Oxygen production with or without nitrite (n=4). **(B)** Oxygen production with varying concentrations of nitrite (n=3). **(C)** Oxygen production with or without ammonium (n=2). Exact cell numbers and raw data can be found in Dataset S1

To test the dependence of oxygen production on NH_4_^+^, cells were inoculated into a near-anoxic medium with 1 mM NO_2_^−^ containing either 2 mM NH_4_^+^ or 0 mM NH_4_^+^, alongside a control with 0 mM NO_2_^-.^ In the presence of NH_4_^+^, oxygen was rapidly consumed to ∼0.3 µM, before production started, as observed previously. Without NH_4_^+^, oxygen consumption was slower, reaching ∼0.3 µM after 1 hour. In both cases oxygen production started immediately after reaching the minimum, with similar production rates in both conditions. After 24 hours, the measured oxygen concentration was slightly higher than conditions with only NO_2_^−^ (4.0 µM with NH_4_^+^ and 4.7 µM without NH_4_^+^), while the variation between replicates was of similar magnitude (Fig. 1c).

### Oxygen production does not convey a competitive advantage under fluctuating oxygen conditions

Recovery after 24 hours in anoxic conditions was tested to determine whether oxygen production would lead to a competitive advantage after anoxic conditions. The same cultures shown in Fig.1a, which after 24 h anoxic incubation had either produced oxygen (with NO_2_^−^) or not (without NO_2_^−^), were returned to optimal oxic growth conditions. Recovery was substantially better for cells that had not produced oxygen than for those that had. Cultures that had produced oxygen showed an extended lag phase of 3 days, after which growth started at slightly lower rates than in the cultures that had not produced oxygen (Fig. 2).

**Figure 2:**
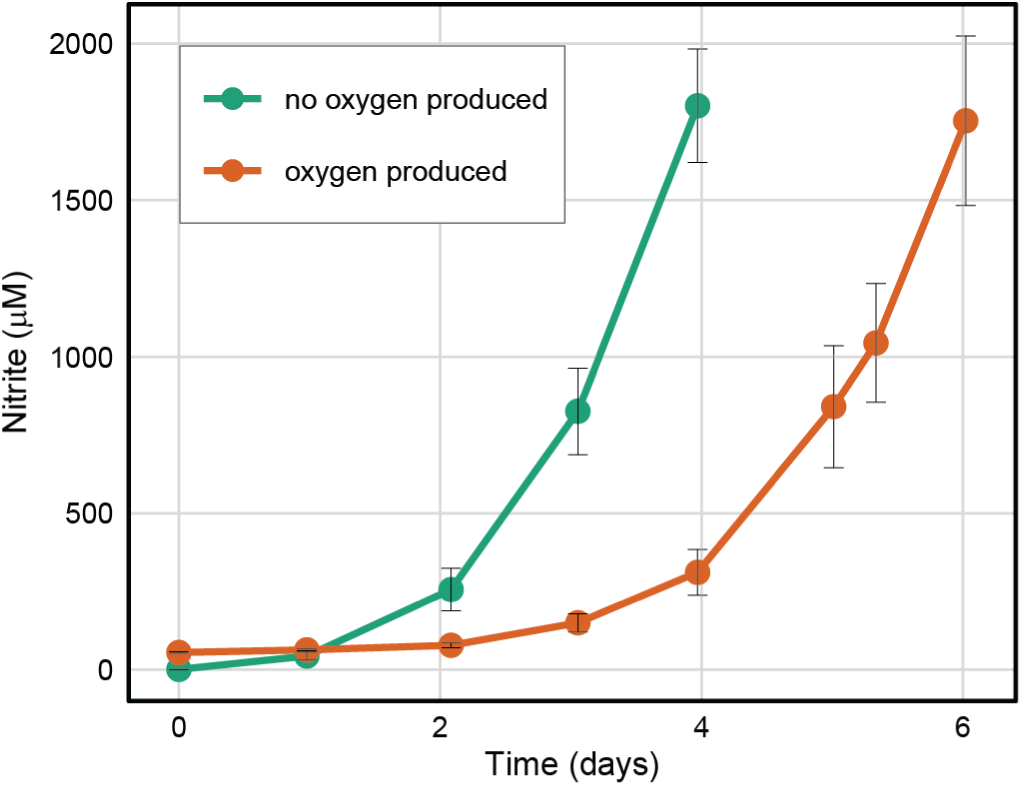
Aerobic recovery after 24h in anoxic conditions. Growth curves of *N. viennensis* in optimal oxic growth medium after either producing or not producing oxygen in anoxia (n=12). Recovery was monitored by nitrite production (µM) over time (days). Exact cell numbers and raw data can be found in Dataset S1.

### Oxygen production is dependent on reducing equivalents

A dependence of oxygen production on NO_2_^−^ would necessitate a supply of electrons from reduced electron carriers (i.e. blue copper proteins) to reduce NO_2_^−^ to O_2_. To test whether the availability of reducing equivalents influences the capacity and amount of oxygen production, cells were starved in normal oxic conditions prior to inoculation into-near anoxic medium containing 1 mM NO_2_^−^ and 2 mM NH_4_^+^. Direct inoculation (no starvation) resulted in the same oxygen consumption and production patterns observed previously. Starvation for 1 or 3 days led to reduced initial oxygen consumption, with a minimum of ∼0.5 µM compared to ∼0.3 µM without starvation (Fig. 3). In all cases, oxygen production started immediately after reaching the respective minimum, but rates were slower in starved cells, i.e. ammonia starvation reduced the capacity for oxygen production. The total amount of oxygen produced was also impacted by starvation. After 24 h, cells without prior starvation produced 4.3 µM oxygen, those starved for 1 day produced 3.6 µM, and cells starved for 3 days produced 2.9 µM (Fig. 3).

**Figure 3:**
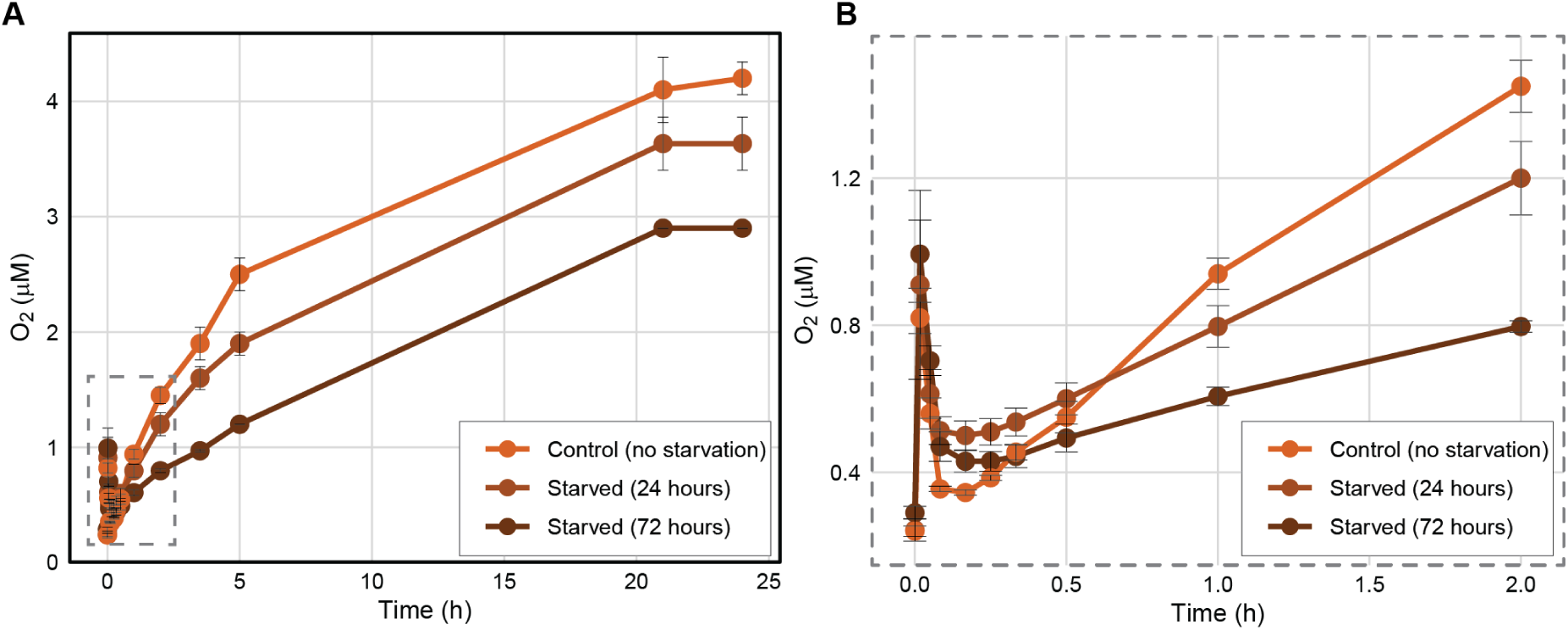
Oxygen production after starvation: Oxygen production (in µM) 0f *N. viennensis* under anoxic conditions over time (in h). All cultures were supplied with 2mM of NH _4_Cl and 1mM NO_2_^−^ after starvation. **(A)** Oxygen production after ammonium starvation in oxic conditions (starved cultures n=3, control n=2). **(B)** A zoomed in view of the dashed boxed in (A) showing the first 2 hours to display different consumption dynamics due to starvation effects. Exact cell numbers and raw data can be found in Dataset S1.

To further test the dependency of oxygen production on the availability of reducing equivalents, the effect of adding oxygen during production in non-starved cultures was tested. An influx of oxygen would allow for the replenishment of reducing equivalents by activating the AMO complex. Cells were inoculated into medium containing 1 mM NO_2_^−^ and 2 mM NH_4_^+^ and divided into two groups. One group received a single oxygen addition of 98.5 µmol (increasing the concentration by 0.86 µM), while the other received three larger additions of 197 µmol each (increasing the concentration by 1.73 µM each time). Additions were made once cultures had produced ∼1 µM oxygen. In both cases, oxygen introductions lead to consumption (Fig. 4a). The consumption of added oxygen was observed even in the case where added oxygen resulted in total oxygen levels below the oxygen production limit of the culture (Fig. 4a). The first two larger additions (increasing concentrations by 1.73 µM) were fully consumed, whereas the smaller addition (increasing concentrations by 0.86 µM) was only partially consumed, reaching a minimum concentration at ∼0.6 µM. When the last 1.73 µM addition was made at 1.5 µM oxygen, consumption again was only partial, reaching a minimum of ∼0.4 µM. Despite these differences in consumption patterns, the final oxygen concentration after 24 h was the same between treatments (Fig. 4). However, total oxygen amounts, calculated from the sum of produced oxygen, were substantially higher in the group with multiple oxygen additions (Fig. 5).

**Figure 4:**
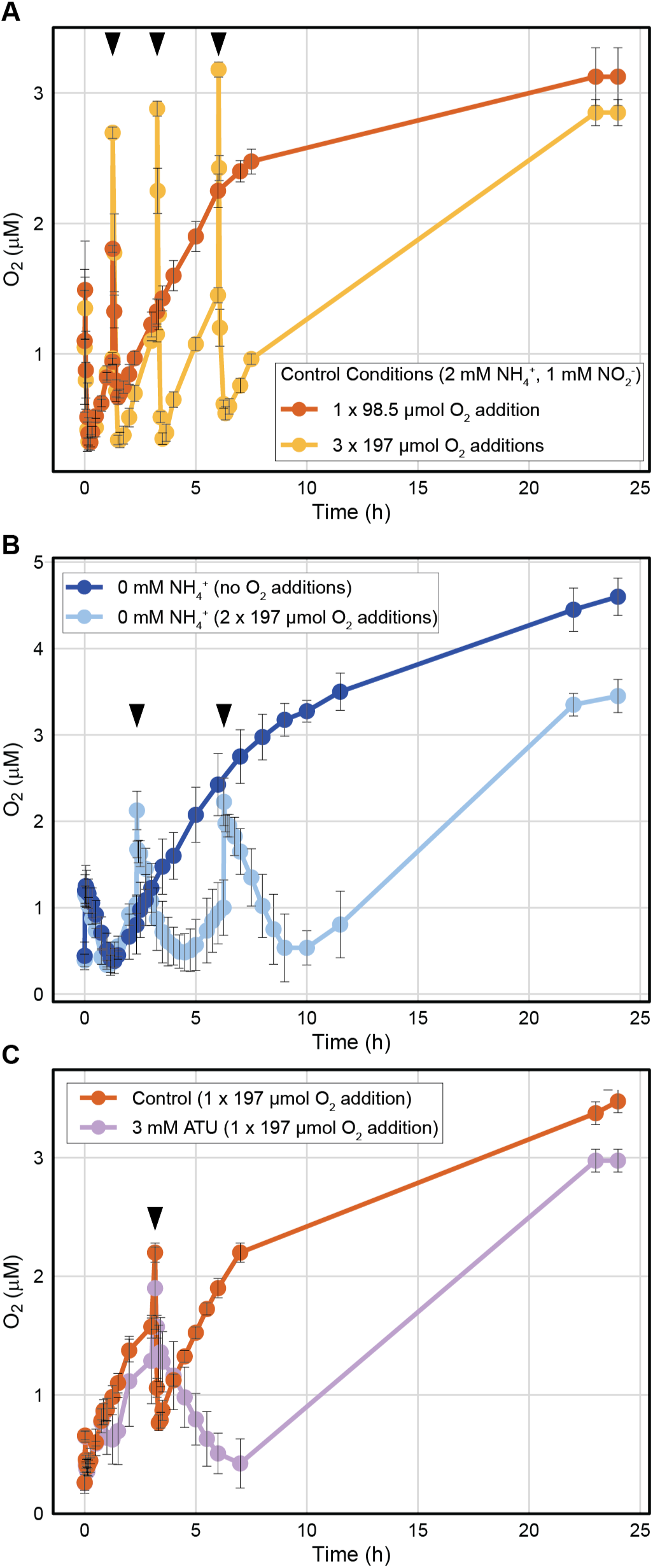
Oxygen production with additions of oxygen: Oxygen production (in µM) under anoxic condition over time (in h). All cultures were supplied with 2mM of NH_4_Cl and 1mM NO_2_^−^ unless specified otherwise. Black arrows indicate time of oxygen addition. **(A)** Production and consumption of oxygen by cultures of *N. viennensis* with the addition of external oxygen(n=4). **(B)** Production and consumption of oxygen with 0 mM NH_4_^+^ by cultures of *N. viennensis* with the addition of external oxygen. (n=4). **(C)** Production and consumption of oxygen in the presence of ATU, a nitrification inhibitor, by cultures of *N. viennensis* with the addition of external oxygen. (n=4).

**Figure 5:**
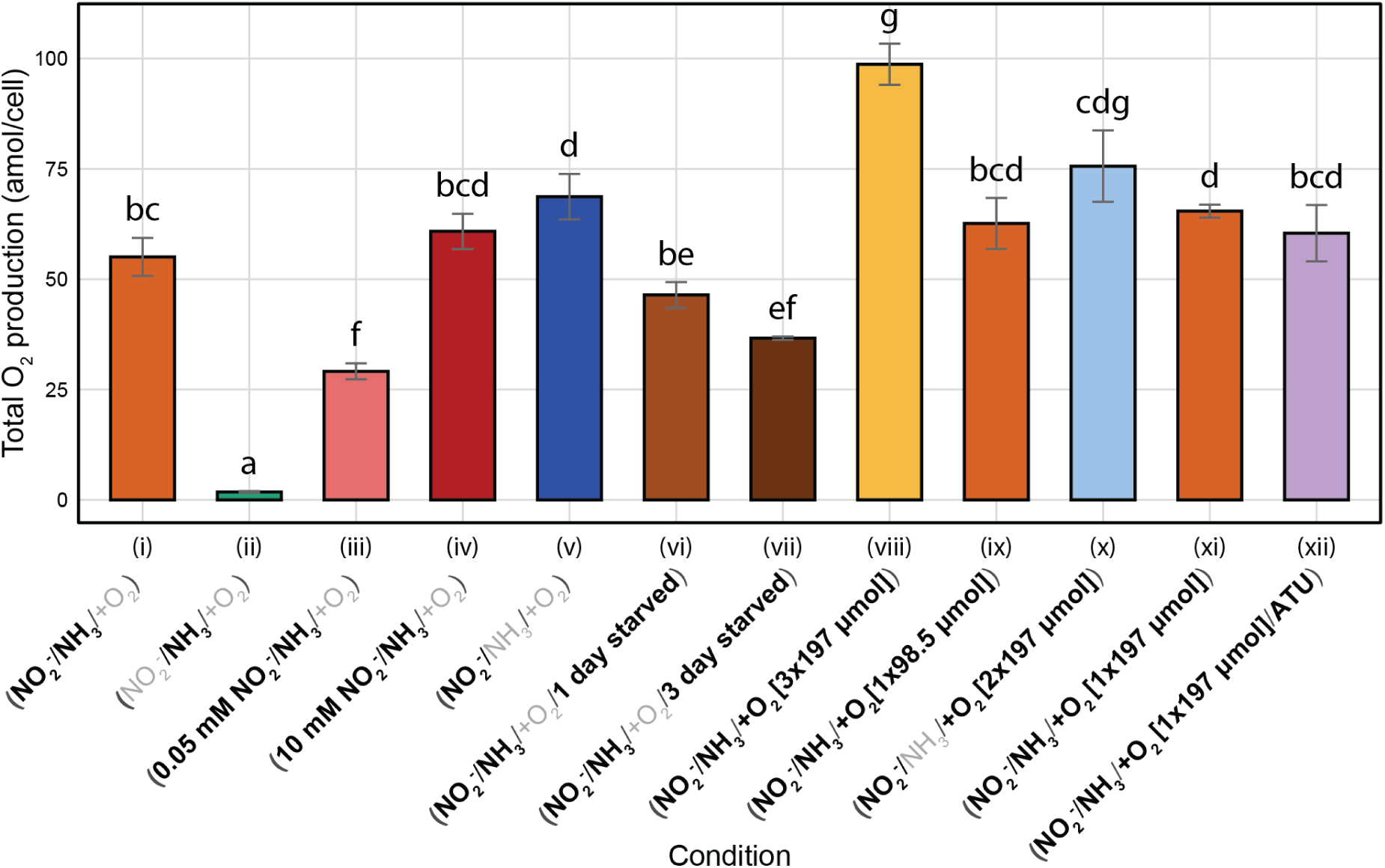
Oxygen accumulation over all experiments: Oxygen accumulation (in amol*cell ^1^) in 24h under different conditions. Figure labels indicate the presence of 2mM NH_4_Cl, NO_2_^−^ (1 mM unless otherwise stated), starvation, ATU, or oxygen additions in bold, or absence in grey. Brackets after oxygen additions indicate the number of additions made and the amount of oxygen introduced by each addition in µmol. Statistical significance between conditions is indicated by letters. Colors of bars correspond to colors in Figures 1-4. Exact cell numbers and raw data can be found in Dataset S1.

The experiment was repeated without NH_4_^+^ in the medium (Fig. 4b). In the absence of ammonia, externally added oxygen should not replenish reducing equivalents. As observed previously (Fig. 1b), initial oxygen consumption was slower, and production began after the minimum concentration of ∼0.3 µM was reached. One group was left undisturbed, while the other received two oxygen additions of 197 µmol (increasing the concentrations by 1.73 µM). Added oxygen was consumed slowly to ∼0.4 µM before production resumed. After 24 h, undisturbed cultures reached an oxygen concentration of 4.5 µM, whereas cultures with additions reached only 3.5 µM. (Fig. 4b) However, the total oxygen production, calculated from the sum of produced oxygen, was remarkably similar (Fig. 5). The experiment was repeated in the presence of NH_4_^+^ but with ATU added to inhibit the AMO complex in the presence of NH_4_^+^ (Fig. 4c). The presence of ATU resulted in slower oxygen consumption and production patterns identical to those observed in cultures without NH_4_^+^ indicating that the observed patterns were due to inactivity of AMO rather than solely an absence of NH_4_^+^.

Figure 5 summarizes the oxygen accumulation normalized to cell number observed across all experiments. Since exact production is unknown, oxygen accumulation is described. Over all experiments, standard oxygen production conditions (1 mM NO_2_^−^ and 2 mM NH_4_^+^) yielded 55±4.1 amol*cell^-1^ (n=9) (Fig 5. (i)). Incubations without NO_2_^−^ did not accumulate significant amounts of oxygen(n=8) (Fig 5. (ii)). Incubations with 0.05 mM NO_2_^−^ yielded a decreased amount of oxygen 29±1.5 amol*cell^-1^(n=3) (Fig 5. (iii)), while incubations with 10mM NO_2_^−^ yielded with 61±3.3 amol*cell^-1^ (n=3) (Fig 5. (iv)). Incubations without NH_4_^+^ led to a higher accumulation of 68.7±4.7 amol*cell^-1^ (n=7) (Fig 5. (v)). When cells were starved, prior to incubations, accumulation was decreased to 46.4±2.4 amol*cell^-1^ (n=3) after one day of starvation and to 36.6±0.3 amol*cell^-1^ (n=3) after 3 days of starvation (Fig 5. (vi, vii)). In the presence of NH_4_^+^, additions totaling 591 µmol oxygen (3 additions of 197 µmol each, increasing the concentration by 1.73 µM each), led to an increased net accumulation of 98.7±4 amol*cell^-1^ (n=4) (Fig 5. (viii)), while a small addition of 98.5µmol (increasing the concentration by 0.86 µM) led to a net accumulation of 62.6±5 amol*cell^-1^ (n=4) (Fig 5. (ix)). In the absence of NH_4_^+^ additions of 394 µmol (2 additions of 197 µmol each, increasing the concentration by 1.73 µM each) of oxygen (Fig 5. (x)) led to an accumulation of 75.6±7 amol*cell^-1^ (n=4). ATU treatment (Fig 5. (xii) led to similar patterns as conditions in the absence of NH_4_^+^ (x), and was similar to the control without ATU (xi), of 60.4±5.5 amol*cell^-1^ (n=4) (Fig 5. (xi, xii)).

### Transcriptomic changes under anoxic conditions

To analyze the change in transcriptomic landscape, transcriptomes were produced after 2 h and 24 h under anoxic conditions in which cells had produced oxygen to a concentration of 1.2 µM or 3.4 µM respectively. After 24 h, the transcriptomic landscape was degraded, leaving only transcripts of the top 30 genes with a TPM >100 and top 250 genes with a TPM >10. 95% of all RNA corresponded to two non-coding RNA (ncRNA) genes: NVIE_RNA1086407D and NVIE_RNA0139656D, both of which are components of RNA-protein complexes (signal recognition particle (SRP) and a ribonucleaseP (RNaseP) respectively).

This is significantly higher than the control for which these transcripts made up 10% of sequenced RNA. The mRNA corresponding to the respective proteins of these complexes was not highly expressed (TPM < 100), and although mRNA and protein levels are not always positively correlated, this led to the assumption that the highly abundant ncRNA genes were not upregulated, but were rather protected from degradation and was the only RNA left in an otherwise degraded transcriptome.

After 2 h, the same trends held true, but to a lesser degree, where 65% of all RNA corresponded to one of the two complexed ncRNAs, while none of the mRNAs corresponding to the associated proteins showed high abundance. Signs of degradation could also be viewed in quality control data of extracted RNA (Supplementary material). Due to the global shifts in transcriptomes based on degradation, a differential expression analysis was not performed.

## Discussion

Ammonia oxidation in AOA relies on oxygen. Oxygen is consumed by AMO and the terminal oxidase, generating a proton motive force that fuels the production of ATP via activity of ATP synthase. Although the specific pathway by which electrons reach the terminal oxidase remains unknown, electrons must also be transferred into the menaquinone pool, which in turn supplies the reducing power required for AMO activity, tightly coupling oxygen consumption, energy conservation, and substrate oxidation (Fig. 6). Thus, all central processes in archaeal ammonia oxidation are intrinsically dependent on oxygen availability.

**Figure 6:**
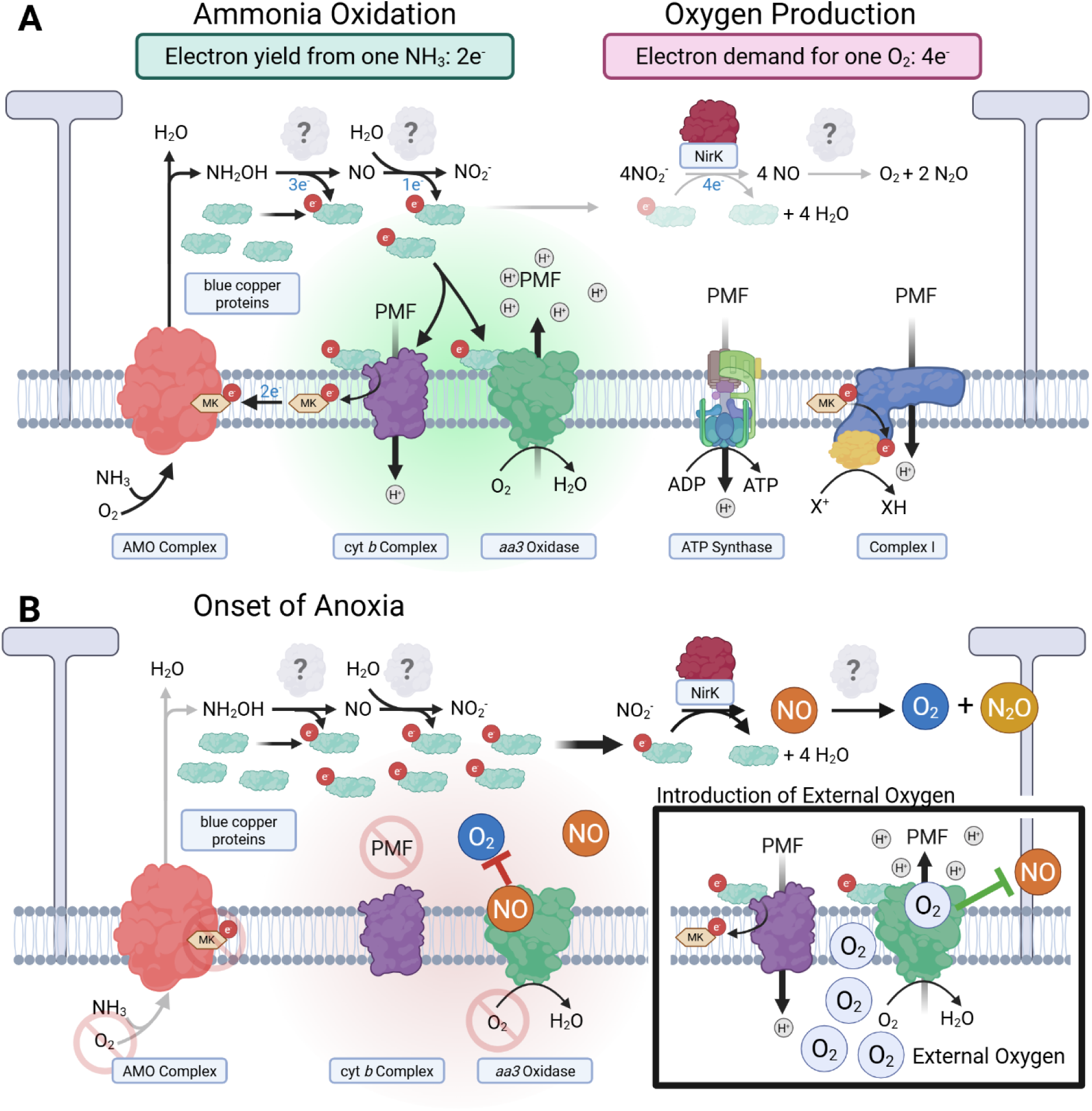
Ammonia oxidation and oxygen accumulation under oxic and anoxic conditions. **A)** Depiction of ammonia oxidation under normal oxic conditions. Electrons extracted from the oxidation of hydroxylamine to nitrite are transferred to blue copper proteins and ultimately are used by the terminal oxidase (*aa3* oxidase) to create a proton motive force (PMF). The produced PMF is used for energy production, the reduction of the quinone pool, and the introduction of electrons into the cell. Oxygen production is only utilized to prevent a build-up of reduced blue copper proteins and reactive intermediates. **B)** Under anoxia the PMF collapses due to lack of oxygen at the terminal oxidase halting all PMF dependent processes and preventing the activation of NH3 by the AMO complex. Reduced blue copper proteins are redirected towards nitrite reductase (NirK) resulting in the production of nitric oxide, oxygen, and nitrous oxide as electron sinks to prevent the build-up of toxic intermediates. Produced oxygen is not utilized due to competitive inhibition at the terminal oxidase with nitric oxide. The black box represents the influence of external oxygen that displaces nitric oxide and allows the cell to function normally until again running out of oxygen. Abbreviations: PMF, proton motive force; MK, menaquinone; NirK, nitrite reductase. Created in BioRender. Hodgskiss, L. (2026) https://BioRender.com/9gb9c5r.

### Oxygen production as an electron overflow pathway

Kraft et al., who first observed oxygen production in marine AOA, suggested that it can sustain ammonia oxidation activity under anoxic conditions, with most of the produced oxygen consumed internally and only a small excess detectable externally ^27^, similar to dark oxygen production in ‘*Candidatus* Methylomirabilis oxyfera’^45^. This was supported by an increase in oxygen accumulation after inhibition of the terminal oxidase by cyanide, i.e., the terminal oxidase was concluded to be actively consuming produced oxygen. Additionally, small levels of N^15^ labeled nitrite were also found to be produced from supplied N^15^ ammonia during oxygen production.

In this study, the ability of AOA to sustain ammonia oxidation while producing oxygen was evaluated by removing NH_4_^+^ from the system to prevent activity of the AMO complex directly. The strikingly similar oxygen production patterns with or without ammonia shown in this manuscript (Fig. 1) makes the simultaneous consumption and production of oxygen unlikely. If oxygen production primarily sustained ammonia oxidation activity under anoxic conditions, incubations without ammonia should have produced substantially more oxygen, or, if a sensing mechanism would be in place, no oxygen. Instead, a slightly higher total production without NH_4_^+^ present in the medium (Fig. 1, Fig. 5), implies a low level/neglectable activity of the ammonia monooxygenase. From the difference in total production (Fig. 5 (i) and (v)), it is estimated, that oxygen production in this setup sustained only minor AMO activity (Dataset S1).

It has also been hypothesized that oxygen production may give an advantage to the cell. However, rather than conferring a physiological benefit, oxygen production impaired recovery from anoxia (Fig. 2). This disadvantage may arise because oxygen production consumes the pool of available reducing equivalents, likely reduced blue copper proteins^13^, to reduce NO_2-_. At the beginning of all experiments with NO_2_^−^, the remaining oxygen is quickly consumed (Fig. 1,3,4). Once depleted, the activity of the terminal oxidase (Complex IV) ceases, collapsing the proton motive force (pmf). Without a pmf, electrons can no longer enter the membrane, halting AMO activity due to a lack of reduced menaquinone. In this scenario, reduced blue copper proteins require an alternative oxidation route.

To address this, it is hypothesized that NO_2_^−^ reduction acts as an electron overflow pathway, alleviating nitrosative stress from NH_2_OH accumulation (that requires oxidized electron acceptors (i.e., blue copper proteins) for the transfer of electrons), while balancing the redox state to maintain efficient ammonia oxidation (Fig. 6). Under oxic conditions, this would be a way to respond quickly to changes in reactive intermediates. In anoxic conditions with reduced blue copper proteins present, this pathway would remain as the sole route for electrons leading to a buildup of the products of electron overflow, mainly NO and oxygen (Fig. 6).

This electron dependent interpretation of oxygen production matches the observations that starvation, which would decrease the pool of reduced blue copper proteins, diminished oxygen production capacities (Fig. 3, Fig. 5 (i) and (vii)). This dynamic effect becomes further evident, when additions of oxygen allow AMO activity to replenish the electron pool in the presence of ammonia (Fig. 4a). In this case, significantly more total oxygen was produced (Fig. 5 (viii)) compared to conditions that had NH_4_^+^ with little (Fig. 5 (ix) or no (Fig. 5 (i)) added oxygen, indicating a limitation that is alleviated by the presence of external oxygen. This limitation may be the availability of reduced blue copper proteins (Fig. 4a) that can be replenished in the presence of both ammonia and externally added oxygen. In contrast, when oxygen additions could not replenish the electron pool (Fig. 4b,c), either because no NH_4_^+^ was available (Fig. 5 (x)) or the AMO was inactivated by ATU (Fig. 5 (xii)), the total production stayed constant compared to control conditions with no oxygen additions and no NH_4_^+^ (Fig. 5 (v)), i.e. the introduction of external oxygen no longer leads to a significant increase in total oxygen production. This is presumably due to an inability of the AMO complex to use external oxygen to replenish the electron pool due to either the absence of NH_4_^+^or inhibition by ATU.

The proposed function as an electron balance pathway parallels the function of nitrifier denitrification in AOB, where nitrite reduction serves as an electron sink from the cytochrome pool during aerobic ammonia oxidation^31,46^, and would also be analogues to heterotrophic denitrification by bacteria or fungi^47,48^. Comparable nitrite based electron sinks have also been described in cyanobacteria^49,50^. NirK has been proposed for several roles in the core metabolism of ammonia oxidizers, either facilitating a reverse reaction of NO to NO_2_^−^ ^11,51^, or as a nitrite reductase, where resulting NO would be involved in the oxidation of NH _2_OH^12^. However, the absence of NirK from the core genome of AOA^52^, especially in the genus *Nitrosocaldus*^53,54^, makes an involvement in core metabolism unlikely. Instead, if NirK functions in an electron overflow pathway, this would imply constitutive expression to rapidly respond to the build up of intermediates. This interpretation is consistent with its constant and high expression levels in many natural environments^51,55–57^, axenic laboratory cultures, and various stress conditions^36,58,59^. Under oxic conditions, oxygen production at these levels would remain undetectable. However, because different AOA strains yield either N_2_O or N_2_, this pathway could act as either a sink or a source of N_2_O^28^, even under oxic conditions. Thus, under oxic conditions this pathway alleviates redox stress from reactive intermediates, but under anoxia it becomes a futile process, accumulating NO and subsequently N_2_O and O_2_. Due to the absence of NirK in the core genome of AOA, it is possible that not all AOA strains utilize this electron overflow pathway and therefore might not produce oxygen in a nitrite dependent manner under anoxia. This opens the possibility that clades missing NirK (i.e., *Nitrosocaldus*) utilize a different mechanism for regulating their electron pool. It is also worth noting that thus far all cultivated strains that have been tested for oxygen production have been isolated from aerobic environments and are strictly dependent on ammonia oxidation coupled to oxygen reduction for growth. It therefore is also possible that specialized lineages in anoxic habitats may have repurposed this unique pathway to sustain activity under anoxic conditions by using other electron donors to fuel this process. This would be in line with the presence of strictly aerobic organisms in permanently anoxic sediments^60^, where dark oxygen production might sustain oxygen dependent metabolism^61^.

### Why is produced oxygen not consumed?

If oxygen production functions as an electron sink during aerobic oxidation, strictly anoxic conditions do not represent the default state of this phenomenon. Nevertheless they yield informative insights. Under anoxia, oxygen production supports only minimal ammonia oxidation (Fig. 1b, Fig. 5), raising the question of why the generated oxygen is not utilized. If this pathway is the sole electron sink under these conditions, even trace amounts of produced oxygen should be consequently consumed via the terminal oxidase and AMO. In contrast to this assumption, oxygen accumulation is observed (Fig. 1), suggesting that the organism cannot use the self produced oxygen. Surprisingly, exogenous oxygen additions trigger consumption of oxygen even when they are supplied below the production threshold (Fig. 4). Additionally, larger oxygen additions resulted in higher consumption (Fig. 4). Once consumption started, even endogenously accumulated oxygen that had previously remained unused, was depleted. Higher oxygen conditions led to a full consumption of oxygen, upon which production started again, while lower additions led to a partial consumption, with production resuming at a concentration determined solely by the amount of oxygen added (Fig. 4). A similar pattern has also been described by Hernández-Magaña and Kraft and was interpreted as being actively regulated by the cell^28^. However, during oxygen production, the transcriptome of *N. viennensis* showed signs of degradation within the first two hours of oxygen production, well before the bulk of oxygen was produced (Dataset S1). This would indicate that the cells were transcriptionally and metabolically impaired, making a robust biological regulation difficult. A regulatory switch based on oxygen concentration also seems unlikely as different behaviors are observed at the same oxygen level; the primary distinction is that consumption is triggered when oxygen is supplied exogenously. If concentration is not the determining factor, an additional element must be involved. This points to the possibility that oxygen consumption is competitively inhibited until a higher amount of oxygen is introduced to displace the inhibitor. Given the central role of the terminal oxidase in oxygen reduction, NO emerges as a compelling candidate.

The current model of oxygen production is dependent on the reduction of NO _2_^−^ to produce NO. Fitting this model, NO accumulation has been observed in all ammonia oxidizers during oxygen production, typically reaching concentrations up to tenfold higher than O ^27,28^. NO can competitively and reversibly bind to the terminal oxidase, inhibiting its activity at nanomolar concentrations in an O_2_ and NO-dependent manner^62,63^. Exogenous O_2_ additions may shift the O_2_:NO ratio in favor of O_2_, displacing NO from the terminal oxidase. Larger O_2_ additions have a stronger effect on this ratio, thereby enhancing O_2_ consumption. Consistent with this mechanism, NO concentrations have been shown to drop following O_2_ additions, after which both NO and O_2_ increased in parallel^28^. The disappearance of NO after exogenous oxygen addition is cohesive with NO itself being an intermediate in the ammonia oxidation process^11,14^; once external oxygen is added and displaces NO, the cell functions as normal until oxygen is again depleted. This offers a plausible explanation for the seemingly counterintuitive lack of a threshold that onsets production or consumption. During undisturbed production, NO and O_2_ accumulate together, resulting in higher, unutilized O_2_ concentrations. The added oxygen however changes this ratio, disassociates inhibition and allows consumption by the terminal oxidase and subsequently also AMO activity, which could alternatively also be directly inhibited by NO. A current unknown in this scenario is the reactivity of NO with the terminal oxidase in AOA. While NO is not reduced in some eukaryotic oxidases, the terminal oxidase of *Thermus thermophilus* has been shown to catalyze the reaction of 2 NO to N_2_O^64^, adding another potential source of N_2_O. While this might account for some of the N_2_O produced in AOA cultures, it would not account for the production of oxygen, or the continued reduction to N_2_ observed in some strains.

### Reconciling oxygen production with ammonia oxidation

Oxygen production has emerged as a shared feature among diverse AOA, yet its physiological role has remained unclear. Because it requires more electrons than ammonia oxidation can provide, it cannot sustain metabolism under anoxic conditions. Instead, the results presented here suggest that oxygen production acts as an electron overflow mechanism, alleviating nitrosative stress and maintaining redox balance during normal growth, similar to nitrifier denitrification in AOB^46^. In line with this, other detoxification mechanisms such as MCO4a have also been proposed^65^, which is not surprising given that organisms with such a highly reactive core metabolism require multiple safeguards. This perspective also resolves several apparent contradictions of the earlier model^27,28,33^: the lack of sustained activity under anoxia, the reduced recovery once electron pools are depleted, and the seemingly wasteful reduction of N_2_O to N_2_, which could act as another electron sink. The high oxygen concentrations observed here under strictly anoxic laboratory conditions therefore represent the product of a detoxification pathway out of its natural, oxic context, and are unlikely to occur in the environment. This fits the observed NH_4_^+^/NO_2_^−^-concentration dependent production of N_2_O mainly, but not exclusively, from NH_3_ under oxic conditions in *N. maritimus*^32^. The precursor for N_2_O would be NO. Fittingly, in this interpretation NO would mainly stem from NH_4_^+^ under oxic conditions with small amounts coming from nitrite when needed to balance electron flow, while under anoxic conditions NO_2_^−^ would be the only source of NO and subsequently N_2_O. Finally, the finding that most self-produced oxygen is not consumed can be explained by the reversible inhibition of the terminal oxidase by NO, which is released once external oxygen is supplied. Taken together, these findings establish oxygen production in AOA not as an adaptation to anoxic conditions, but as a redox balancing strategy that safeguards cells under nitrosative stress.

## Acknowledgements

We thank Dr. Maximilian Dreer and Dr. Melina Kerou for discussions. The RNA sequencing was performed by the Next Generation Sequencing Facility at Vienna BioCenter Core Facilities (VBCF), member of the Vienna BioCenter (VBC), Austria. The computational results of this work have been achieved using the Life Science Compute Cluster (LiSC) of the University of Vienna. This project was funded by the Austrian Science Fund, Projects P36287 (The Ammonia Oxidation Process in Archaea) and Z437 (Archaea Ecology and Evolution), as well as EU Horizon 2020 twinning project ActionR (Research Action Network for Reducing Reactive Nitrogen Losses from Agricultural Ecosystems) No. 101079299.

