## Supplementary_Material for "Oxygen production as an electron overflow pathway in ammonia-oxidizing archaea"

**This PDF file includes:**

Figs. S1 to S4

**Other Supplementary Materials for this manuscript include the following:**

Dataset S1 – Associated oxygen production and growth data for Figures 1-5.

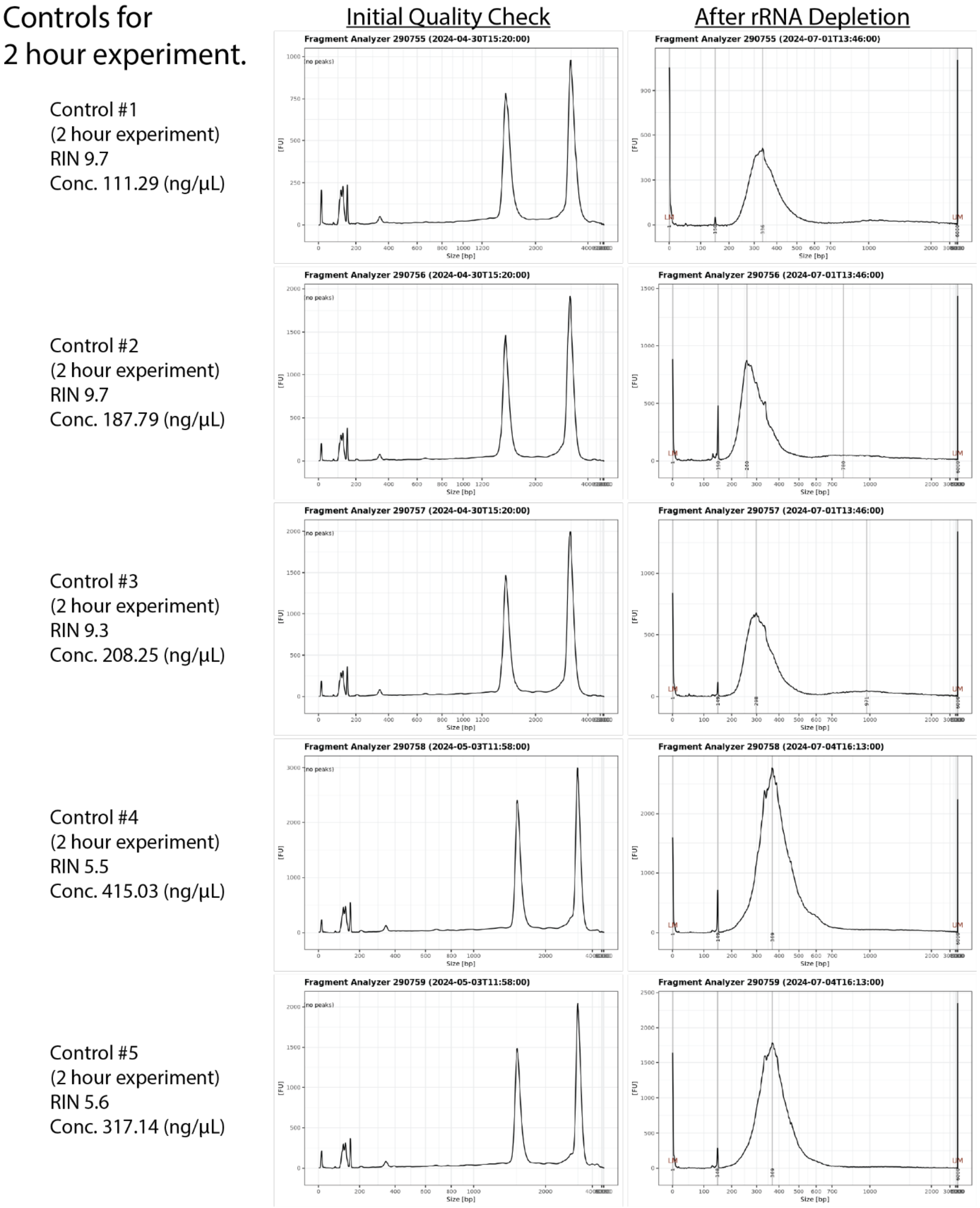


**Fig. S1.**

Bioanalyzer data for controls of 2 hour anoxia experiment. Abbreviations: RIN, RNA integrity number.

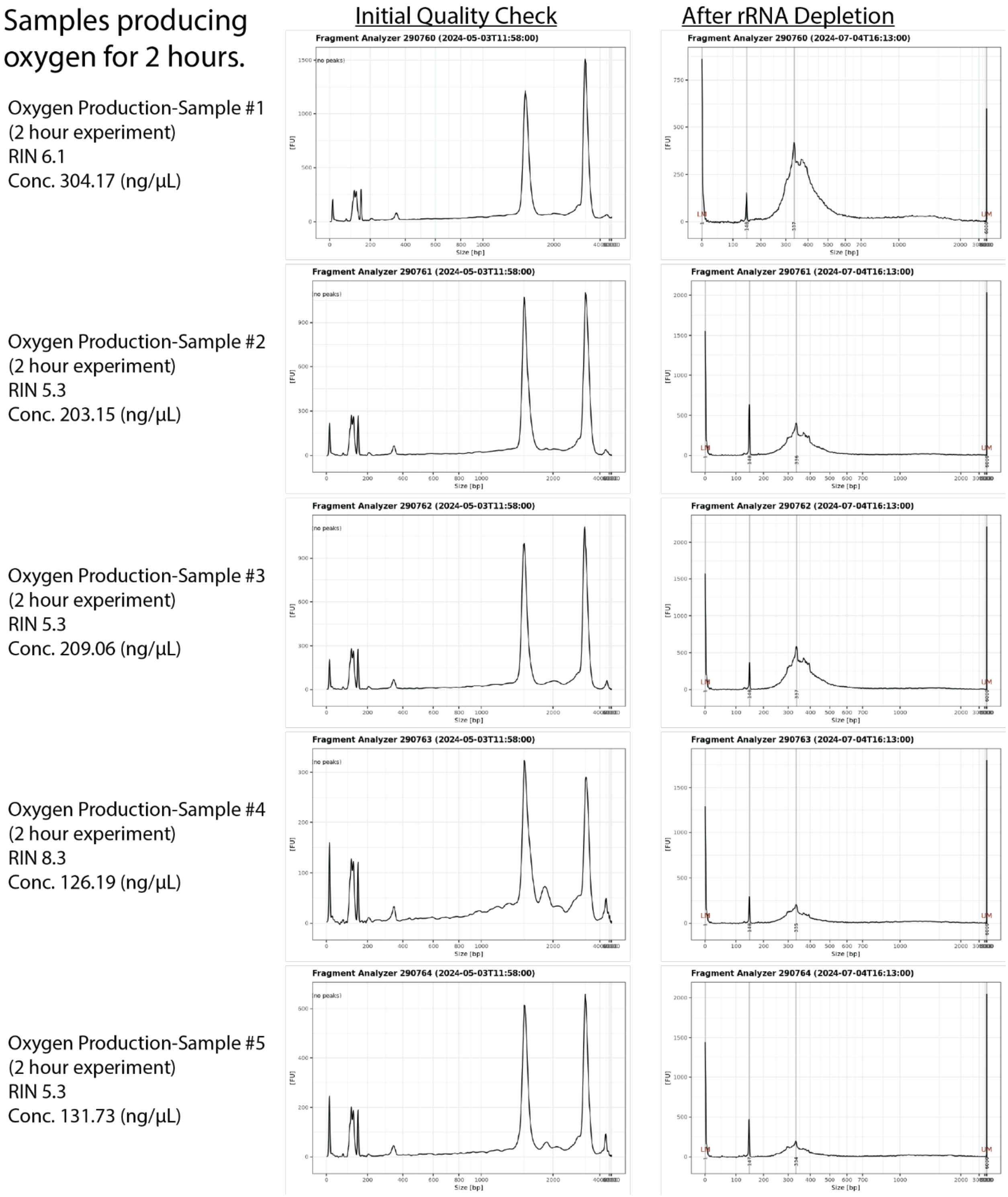


**Fig. S2.**

Bioanalyzer data for samples of 2 hour anoxia experiment. Abbreviations: RIN, RNA integrity number.

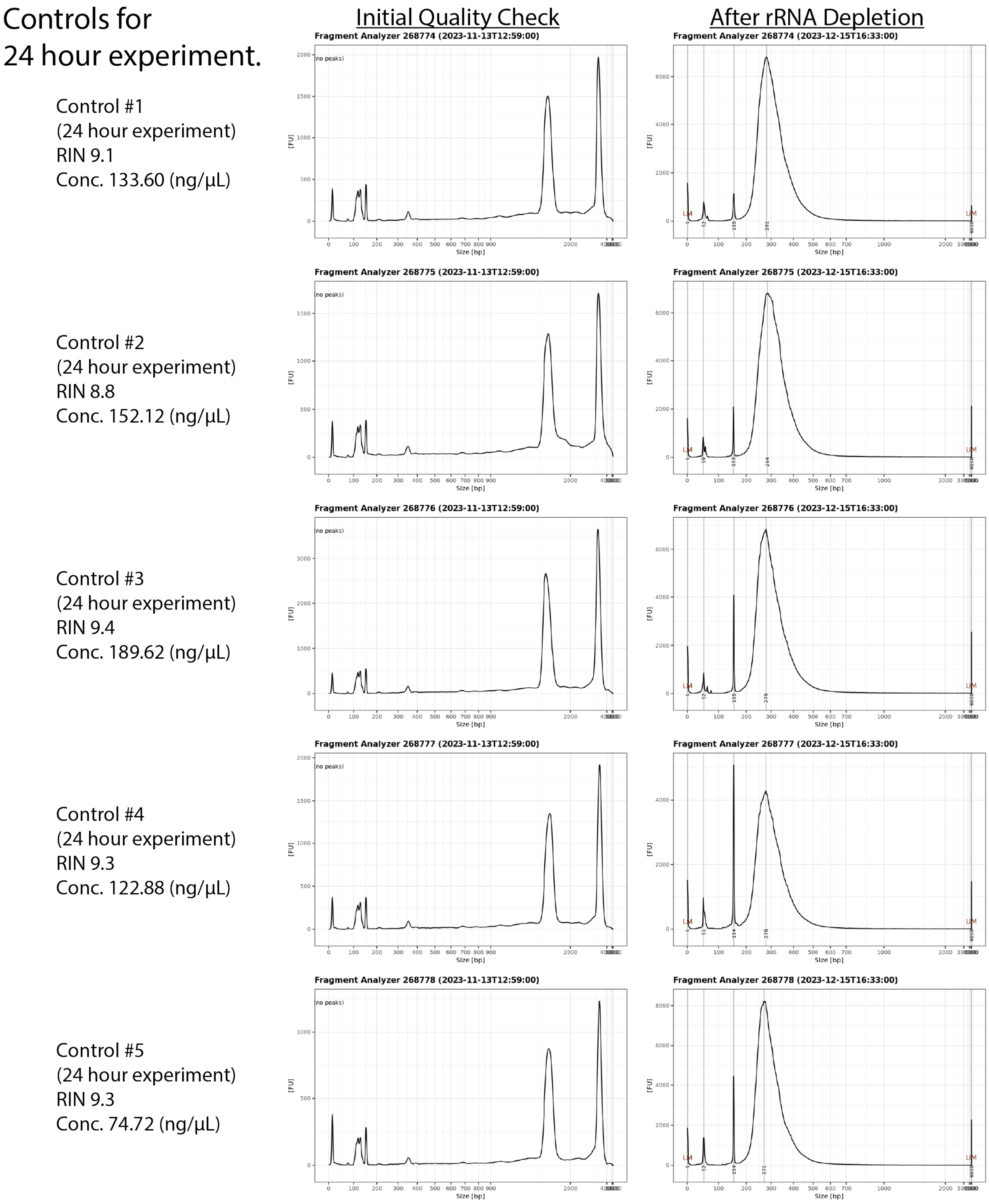


**Fig. S3.**

Bioanalyzer data for controls of 24 hour anoxia experiment. Abbreviations: RIN, RNA integrity number.

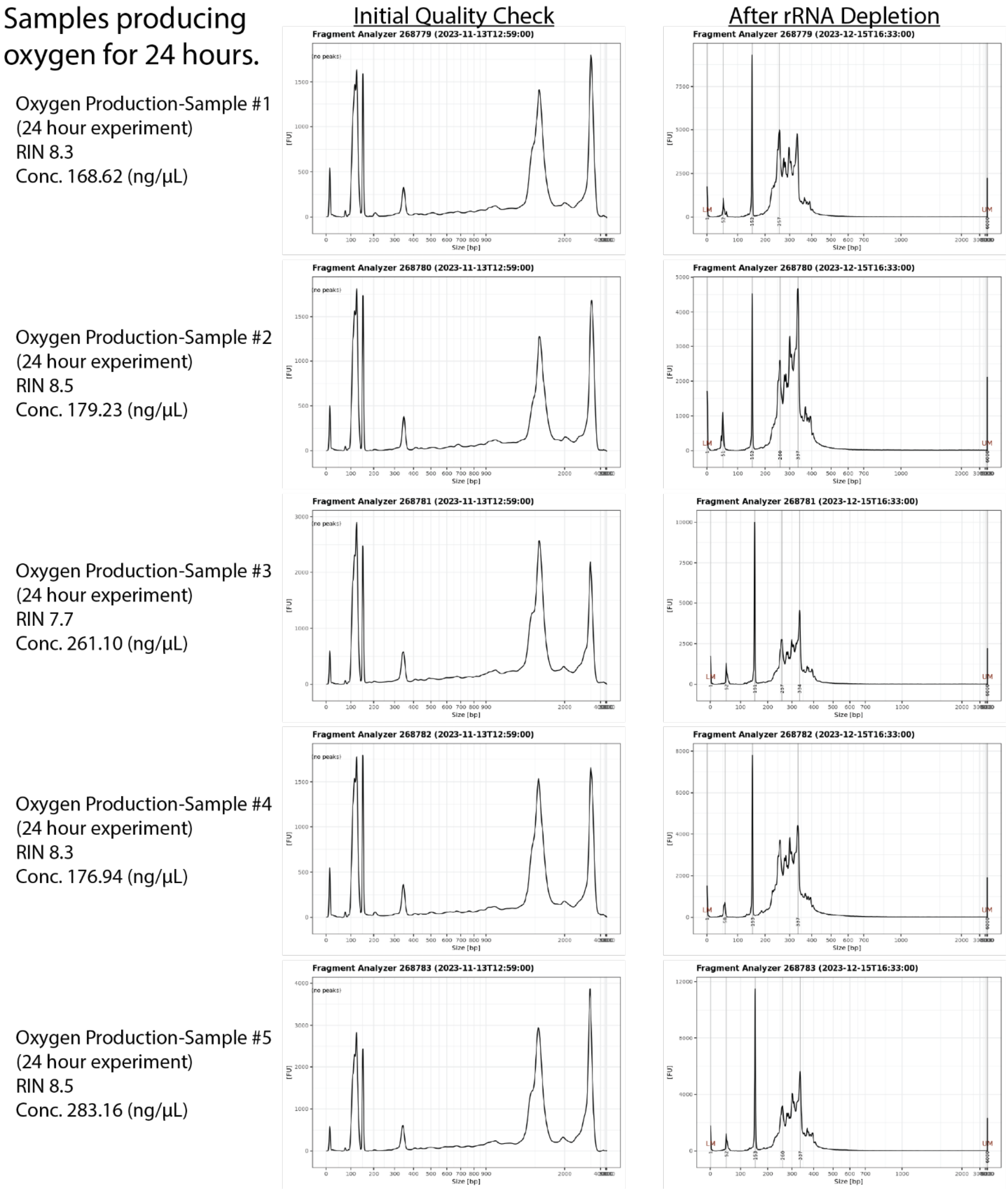


**Fig. S4.**

Bioanalyzer data for samples of 24 hour anoxia experiment. Abbreviations: RIN, RNA integrity number.
